# β-lactoglobulin a new whey: Computational redesign improves stability and nutritional composition

**DOI:** 10.64898/2026.08.17.745312

**Authors:** Maija Greis, Ulysse Castet, Elsa Berlin, Sofia Klangby, Oliwia Bancerz-Aleksiejczuk, Francisco Vilaplana, Julia K. Keppler, Elton P. Hudson

## Abstract

Protein engineering and precision fermentation provide an opportunity to increase the value of food proteins by improving their solubility, stability, functionality, or nutritional composition. Here, we use β-lactoglobulin (βLG) as a model protein to investigate how state-of-the-art computational protein design approaches affect these properties. First, the deep learning-based design tool ProteinMPNN was used to alter up to 20% of βLG residues for increased stability. Second, the physics-based modeling platform PyRosetta was used to find positions in βLG accommodating increased branched-chain amino acid (BCAA) content and up to 10 residues were simultaneously exchanged. Experimental characterisation of ProteinMPNN and stabilised BCAA-enriched variants showed similar secondary structure and oligomeric state as native βLG. ProteinMPNN variants gave increased titers and increased thermal stability up to 15 °C, and this correlated with changes in the rate of surface pressure in droplet tensiometry. Stabilized BCAA-enriched mutants had altered acid solubility. Correlations between computationally derived biophysical metrics and experimental properties are presented and suggest some predictive power for surface hydrophobicity on protein yield.

## Introduction

Artificial intelligence (AI) is revolutionising food innovation by enabling predictive and design-driven food development^1^. One of the largest impact areas is to channel current state-of-the-art protein design approaches to develop food proteins with increased functional properties. To date, the majority of successful biotechnical applications of protein design have been in enzyme engineering^2^. These tools are largely optimized to improve general protein characteristics such as structural stability. Protein structure prediction tools have been used successfully to improve industrial enzymes such as lipases, making them work more efficiently also in the food industry^3,4^. In addition to enzymes, natural sweet proteins are attractive targets for protein engineering because they combine a well-defined receptor-binding function with challenges such as poor thermal stability, enabling targeted optimization of specific properties. For example, thermal stability has been improved for sweet protein Neoculin^5^ and Monellin^4^, both by PyRosetta. In another proof-of-concept study, protein design was used to enhance the thickening properties of potato protein patatin by introducing surface cysteines, which also increased its sulfur amino acid content^6^. This work demonstrated that computational protein engineering can modulate food-relevant properties such as thermal stability and gelation. Examples of completely *de novo* designed proteins for food applications remain scarce. However, recent work has explored the use of generative protein language models to design de novo proteins toward a target amino acid composition, highlighting nutritional composition as a potential objective for computational protein design^7^.

Food proteins can function in various foods by an extension of their biological function. The majority of food proteins perform multiple functions that depend not only on stability, but also on conformational flexibility, intermolecular interactions and surface properties. Food functionality emerges from multiple structural and physicochemical properties^8^. Unlike many biological proteins, food proteins often perform their technological functions after partial unfolding rather than in their fully native state^9^. Thus, increasing protein stability may also influence food-functional properties that depend on these structural transitions. Protein engineering provides an opportunity to investigate how changes in protein structure affect these food-functional properties. The extent to which optimization for stability translates into desirable food-relevant properties remains unclear.

Another important aspect of protein quality is the amount and digestibility of essential amino acids (EAAs), which cannot be synthesized in sufficient amounts by the human body and must therefore be obtained from the diet^10^. Three of the nine EAAs, leucine, isoleucine and valine, are branched-chain amino acids (BCAAs)^11^. BCAAs have an important role in protein metabolism, and leucine in particular stimulates muscle protein synthesis^12, 13^. Although BCAAs can stimulate muscle protein synthesis, it has been shown that a complete protein containing all EAAs produces a greater response^11^. Protein design could therefore be used to increase the amount of selected nutritionally valuable amino acids within a complete protein. This could provide more of these amino acids per gram of protein, which may be useful in foods where only a limited amount of protein can be added.

One protein of interest is β-lactoglobulin (βLG), a milk protein consisting of 162 amino acids (18.3 kDa). βLG is a widely studied whey protein and a useful model for engineering next-generation food proteins. It can be produced recombinantly in *E. coli*^14, 15^ and at larger scale in yeast expression systems. βLG exhibits valuable techno-functional properties, including foaming and emulsification through interfacial film formation^16^, and heat-induced gelation driven by partial unfolding and disulfide-mediated aggregation^17^. Previous protein engineering of βLG has mainly focused on targeted mutagenesis to investigate structure–function relationships. Mutations have been shown to alter properties such as dimer formation, aggregation behavior, and gel strength^18,19,20^. Hoppenreijs et al. further showed that changes in cysteine residues, together with differences between natural βLG isoforms, can substantially affect its foaming and gelation properties^21^. Even one or a few amino acid substitutions can result in clear difference in βLG thermal stability. This has been reported between natural variants and following targeted cysteine substitutions^22^. However, most studies have examined individual properties rather than the simultaneous optimization of multiple functional traits.

Thermal instability is a challenge for the dairy industry as denaturation of βLG causes off-flavours and aggregation causes fouling of equipment^23^. In addition, protein engineering could provide an opportunity to produce βLG with an increased BCAA content, enabling greater nutritional impact with smaller amounts of protein^11^. Protein engineering using computational methods is a very active field of research, but their application to major functional food proteins, such as βLG, remains limited.

ProteinMPNN redesigns amino acid sequences by learning sequence–structure relationships from experimentally determined protein structures^24^. Although the model is not explicitly trained to optimize thermal stability, the generated sequences often exhibit improved folding and thermal stability by increasing compatibility with the target protein backbone. By redesigning amino acid sequences while preserving the protein backbone, ProteinMPNN has enabled the development of proteins with enhanced thermal stability^2^.

Food functionality is controlled by multiple interacting molecular properties, but it is still poorly understood how these properties translate into functionality. Protein engineering combined with computational and experimental characterization provides a way to investigate these properties. Here, we use βLG as a model system to explore two questions: can its BCAA content be increased through protein design while maintaining its solubility and stability, and can thermal stability and solubility be enhanced while preserving food-relevant functionality?

## Results

### Computational redesign of bovine βLG

We applied two orthogonal computational design approaches to βLG: Generative deep-learning sequence recovery using ProteinMPNN and a physics-guided mutation using PyRosetta **(Fig. 1)**. For both approaches, βLG-B was used as a reference wild type and starting sequence. Variant B was selected based on its higher denaturation temp compared to A^17,22^ and its characterization in previous studies.

**Figure 1.**
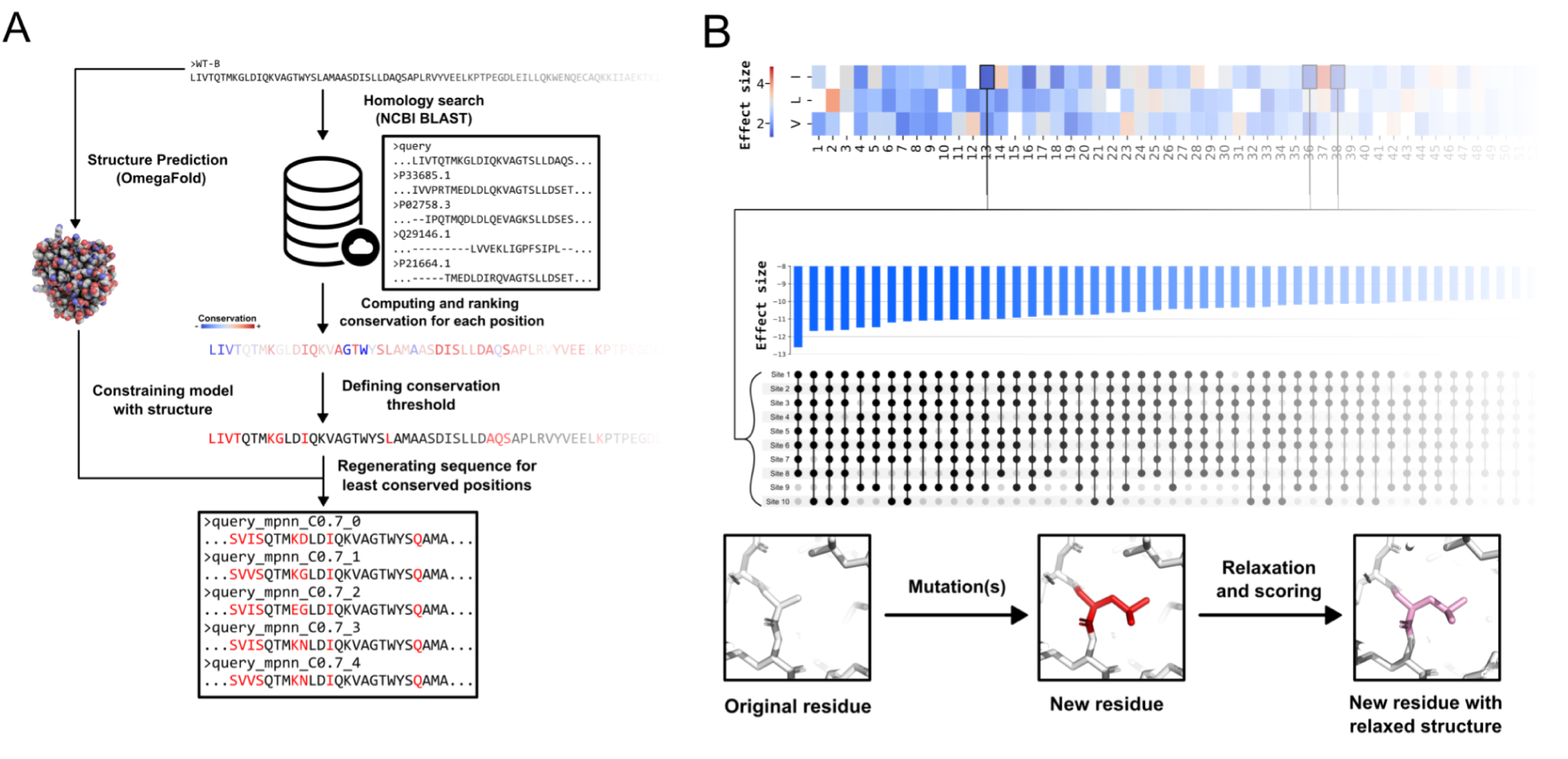
Design workflow for computational generation of candidate variants. (**A**) Generative deep-learning sequence recovery using homology-constrained ProteinMPNN. An MSA is built from the starting WT-B sequence and used to quantify residue conservation. ProteinMPNN is provided with the index of the x least conserved positions, along with the structure of starting sequence, to generate new sequences differing specifically at the selected sites. (**B**) Physics-guided structure editing using PyRosetta. Mutational scan of all possible single Branched-chain amino acid (BCAA) substitutions (top). Selection of the top 10 best scoring mutations. Exhaustive scoring of all possible subsets (middle upset plot). For a specific mutant, harboring one or more mutations, 50 Ref2015 scores are computed and averaged after sequential application of the mutations and downstream relaxation using FastRelax (bottom).

For ProteinMPNN, we allowed variation of up to 20% of the protein sequence. To restrict which residues the model could alter, we first built a multiple sequence alignment (MSA) of 71 βLG homolog sequences to rank the degree of conservation of residues across βLG evolution, in a workflow similar to Sumida et al.^2^ (**Fig. 1A and Supp. Data**). Residues that are most conserved in evolution are assumed to be critical for protein fitness stability and function, and therefore less tolerant for mutation. Next, nested sets of residues were selected based on three different conservation thresholds: 95%, 90%, and 80%. These sets were used as masks to limit ProteinMPNN redesign to only residues below the conservation threshold. Sequence generation was constrained to the 3D structure of βLG-B, as predicted with OmegaFold^25^. A total of 2858 sequences were generated. To guide selection of candidates for experimental testing, a set of sequence-and structure-based quality metrics were computed for each sequence. These included sequence identity, root mean square deviation (RMSD) compared to βLG structure, and an average predicted local distance difference test (pLDDT). We then selected 13 ProteinMPNN variants for synthesis and purification, four from conservation cutoffs 80% (MPNN_80_1-4) and 90% (MPNN_90_1-4), and five from conservation cutoff 95% (MPNN_95_1-5; **Table S1**). Each variant selected for expression had a pLDDT higher than 90, indicating high confidence in structure prediction, and an RMSD < 2 Å, indicating a predicted secondary structure similar to βLG-B.

In a second approach, we used functions from the PyRosetta toolkit^26^ to estimate protein fitness tolerance to residue substitutions (**Fig. 1B**). Considering the importance of branched-chain amino acids (BCAA; leucine, isoleucine, and valine) in the nutritional properties of βLG^11^, our goal was to create βLG variants with increased BCAA content which retained stability. The starting structure for PyRosetta was an AlphaFold prediction of βLG variant B^27^. The relative stability of each variant (ΔΔG) was estimated using the Ref2015 energy function^28^, which has previously been shown to be a reasonable predictor of protein stability changes upon mutation. We performed a full computation mutational scan, substituting each residue with either L, V, or I (445 variants). The predicted impact of mutation on protein fitness was estimated using Cohen’s d effect size between the unmutated and post-mutation scores (**Supplementary Fig. 3)**.

The mutational scan results included some notable features. Mutations of small residues, such as alanine or glycine, were predicted to be detrimental to protein stability, likely illustrating the resulting steric clashes arising from BCAA substitutions. On the contrary, all mutations of E89 showed a strong stabilizing effect on the protein compared to other positions. E89 is particularly relevant for ligand binding because it acts as a pH-sensitive switch controlling opening of the EF loop and access to the central calyx^29^. Thus, substitution of E89 may alter the normal pH-dependent accessibility of the ligand-binding site. This would suggest a potential trade-off between stability and ligand binding function at this residue.

To generate multi-site BCAA variants, we extracted the 10 non-overlapping BCAA mutations predicted to have the most positive impact on protein stability. All of these substitutions were then independently evaluated (1013 variants). A similar combinational approach was also performed using the 10 non-overlapping BCAA mutations predicted to have the most neutral impact on protein stability (i.e. ΔΔG closest to zero). From these multi-site variants, we selected 10 for experimental testing: the most stabilized variants containing either 4, 6, 8, and 10 substitutions (BCAA_stab_1-4; Cohen’s *d* −12.59 to −9.29), the most neutral variants containing either 4, 6, 8, and 10 substitutions (predicted stability change closest to zero; BCAA_neut_1-4), and two single-site substitution variants predicted to be significantly de-stabilized (BCAA_neg_1-2; Cohen’s *d* = 7.9 and 9.7). We note that βLG-B contains 41 BCAA; so a 10-point mutant increases this to 51 BCAA, a 25% increase in BCAA content.

For each mutant, we computationally estimated pI, molecular weight, aromaticity, and secondary-structure-related quantities (**Table S2**). We also calculated sequence distance among variants, BCAA variants were highly similar to each other, as expected as they were constructed by adding sequential substitutions. ProteinMPNN variants were significantly different, even within the same conservation group. For example, variant MPNN_80_1 and MPNN_80_2 different from βLG-B at 26 and 28 positions, respectively, and from each other at 11 positions (**Fig. S2**).

### βLG mutants retain secondary structure and exist as dimers

A first round of characterization was performed for five natural βLG variants and all 13 ProteinMPNN variants **(Table S1)**. Proteins were produced in *E. coli,* purified, and assayed for thermal denaturation. Based on these results, βLG-B and six ProteinMPNN-designed variants, two from each conservation cutoff, were selected for further characterisation **(Table 1)**. These variants were re-expressed and purified using affinity chromatography with subsequent removal of affinity tags. The protein yield exceeded that of WT-B **(Fig. 2A; Table S3).**

**Figure 2.**
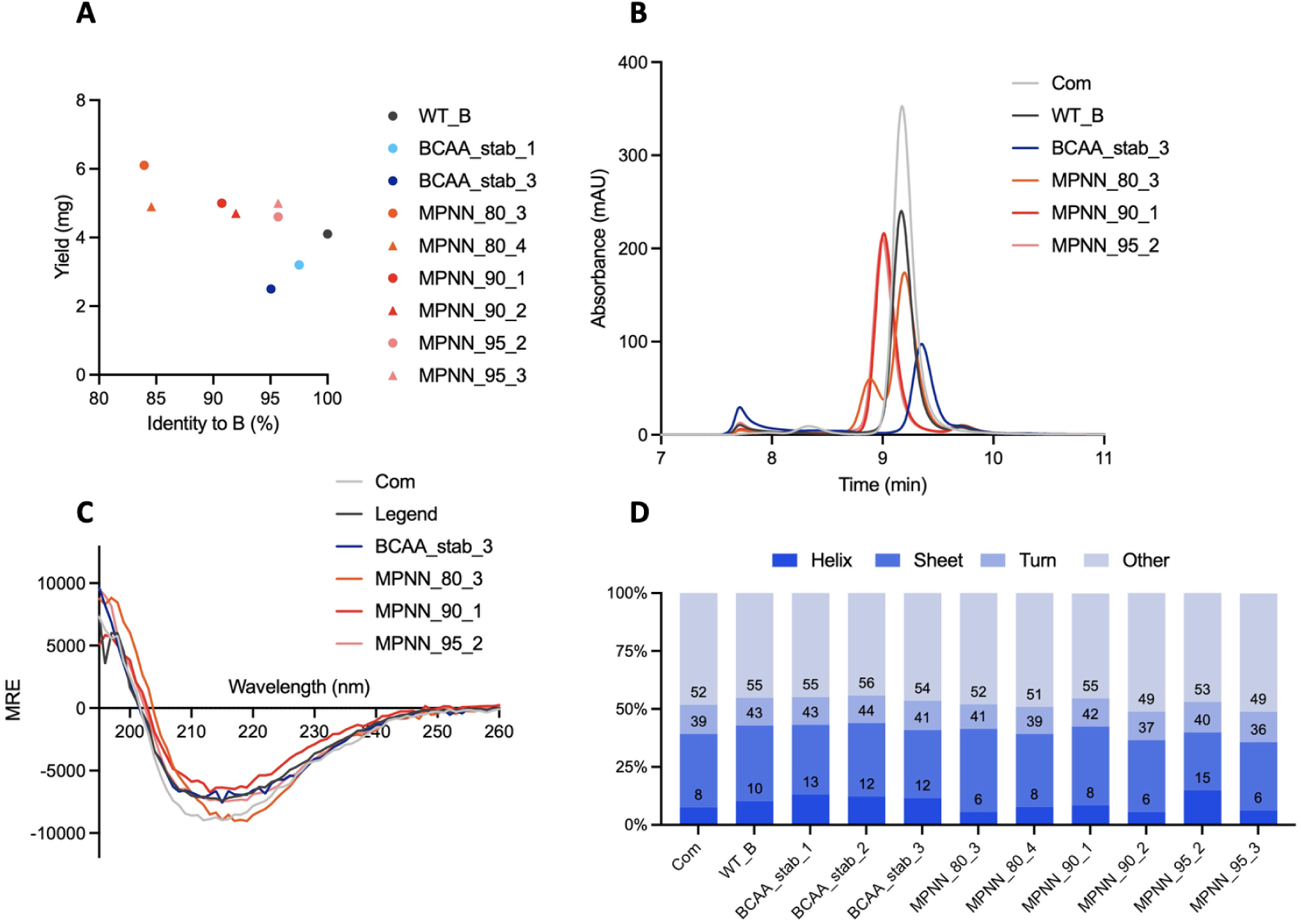
Expression and structural characterization of β-lactoglobulin variants. **(A)** Protein yield as a function of sequence identity to WT_B. ProteinMPNN-designed variants, and five BCAA-enriched variants. (B) Size-exclusion chromatography (SEC) profiles of selected variants. (C) Far-UV circular dichroism (CD) spectra of selected variants. (Signal reported in molar residue ellipticity (MRE)). (D) Secondary-structure composition estimated from CD spectra using BeStSel.

**Table 1.** A total of 10 variants, WT_B, six proteinMPNN, and three BCAA variants were chosen for the detailed characterization.

| Variant name | pLDDT | rmsd_wtB | nb BCAAs |
| --- | --- | --- | --- |
| βLG-B | 95.569 | 0 | 41.000 |
| MPNN_80_3 | 94.592 | 1.17 | 42.000 |
| MPNN_80_4 | 92.085 | 1.44 | 42.000 |
| MPNN_90_1 | 91.766 | 1.13 | 38.000 |
| MPNN_90_2 | 93.317 | 1.23 | 40.000 |
| MPNN_95_2 | 1.616 | 0.43 | 38.000 |
| MPNN_95_3 | 1.606 | 0.45 | 39.000 |
| BCAA_stab_1 | 1.604 | 0.36 | 45.000 |
| BCAA_stab_2 | 1.645 | 0.5 | 47.000 |
| BCAA_stab_3 | 1.641 | 0.4 | 49.000 |

In parallel, the ten BCAA-enriched variants designed using PyRosetta were cloned and expressed in *E. coli*. BCAA_neg_1 and BCAA_neg _ 2, containing the single substitutions N88I and G17I, respectively, did not yield detectable soluble protein. Of the BCAA_neutral variants, only BCAA_neut_1 and BCAA_neut_3 produced soluble protein but yields were insufficient for further characterization. Of the BCAA stabilised variants, only BCAA_stab_1-3 gave high soluble protein yields and were successfully purified without affinity tags for further characterisation (**Fig. 2A)**.

The purity of all recombinant βLG variants was estimated as >90% by HPLC-SEC (**Fig 2B)**, similar to that reported previously for recombinant βLG produced in *E.coli*^15^. Fractions corresponding to proteins above 60 kDa and residual tagged βLG were minor compared with the predominant monomeric and dimeric βLG fractions. All samples showed a weak band at approximately 90 kDa by SDS-PAGE (**Fig. S3**), while variants MPNN variants 80_3 and 80_4 contained traces of βLG retaining the affinity tag. Purified variants were also characterized by MALDI-TOF mass spectrometry, and all showed the expected molecular weight (18 kDa monomer).

No masses corresponding to tagged βLG or higher-molecular-weight proteins were detected by MALDI-TOF mass spectrometry **(Table S3**).

The secondary structure of purified variants was assessed using circular dichroism spectroscopy (CD). The spectra of all variants closely resembled that of the commercial reference (**Fig. 2C)**. βLG-A and βLG-B have previously been reported to have a CD minimum at 216 nm at pH 7.4^30^, which is close to the minimum at approximately 215 nm observed here under similar conditions. Content of α-helices and 6–15% and β-sheet, as calculated from CD spectra, ranged from 6-15% and 36–44%, respectively **(Fig. 2D; Table S5**).

Ruminant βLG is mainly dimeric at physiological pH but tends to monomerize at low pH, low ionic strength^31^. The commercial reference showed a single peak corresponding to βLG dimer, with no detectable monomeric species. The oligomeric state of most βLG variants was >95% dimer, as analyzed by HPLC-SEC (**Table S4**). BCAA_stab_3 was the only exception, showing an increased monomer fraction (10%) and larger elution fractions above 60 kDa compared with other variants.

### βLG mutants have enhanced thermal stability

The commercial reference and purified βLG-B had melting temperatures of 72.5 °C and 73.8 °C, respectively as measured by differential scanning fluorimetry (nano-DSF), consistent with previous reports. The melting temperatures for all ProteinMPNN variants were higher, ranging from 74.4 °C to 88.3 °C, and the melting temperature increased with the extent of mutation (**Fig. 3B, Table S6**). BCAA variants were marginally stabilized (BCAA_stab_1 and BCAA_stab_3 with T_m_ 77.8 °C and 73.5 °C, respectively). NanoDSF relies on intrinsic tryptophan fluorescence, which is highly sensitive to changes in the local environment^32^. Thus, amino acid substitutions may affect the fluorescence profiles independently of protein unfolding.

**Figure 3.**
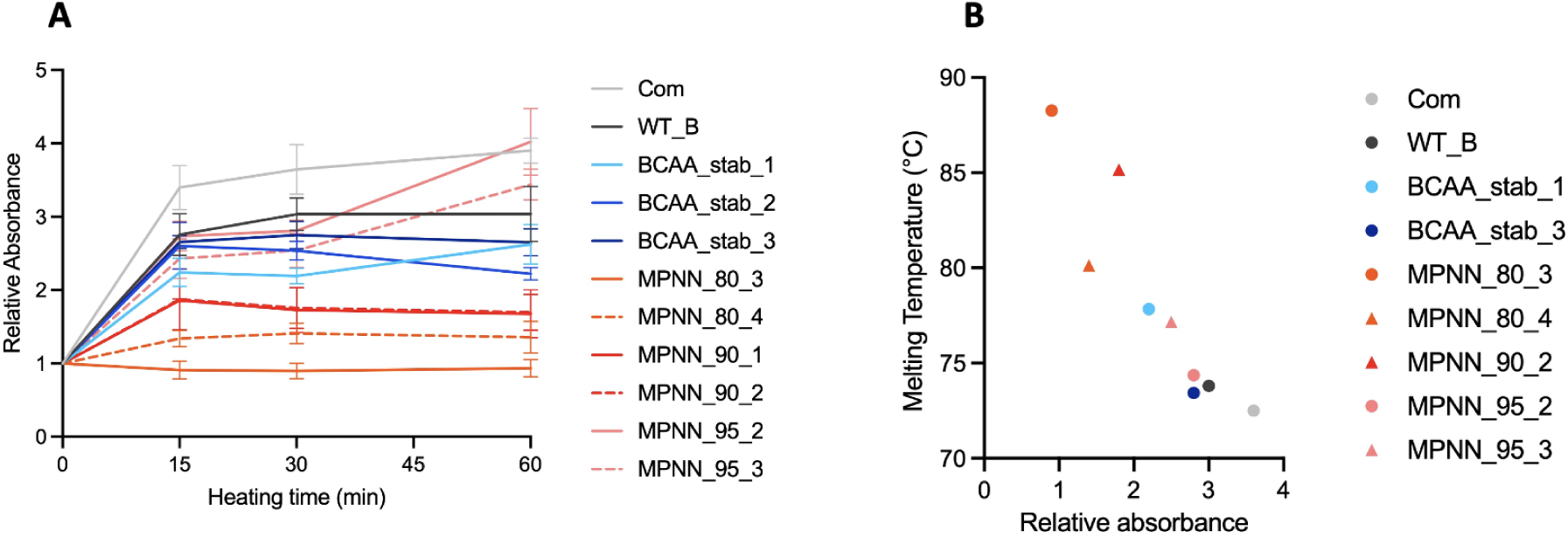
Thermal stability and free thiol content of β-lactoglobulin variants. **(A)** Free thiol content during incubation at 70 °C for 60 min, determined using Ellman’s assay, for the commercial reference and selected ProteinMPNN-designed from each sequence-identity group and BCAA-enriched variants **(B)** Higher melting temperatures correlate with lower free thiol levels (ρ = −0.934, *p* < 0.001).

Thermal stability was also assessed using changes in absorbance of Ellman’s reagent, which measures exposure of thiol groups upon heat-induced denaturation. ProteinMPNN variants maintained comparatively low absorbance responses over the heat treatment, apart from variant MPNN_95_2 which had similar absorbance to the commercial βLG reference **(Fig. 3A)**. The BCAA variants showed absorbance profiles closer to those of βLG-B and the commercial reference, although their absorbance remained slightly lower throughout the heating period, indicating higher thermal stability. There was a strong correlation between protein unfolding as measured by Ellman’s reagent and by nanoDSF (**Fig. 3B).**

### Altered aggregation propensity and surface absorption in some βLG variants

Acid solubility was assessed as an indirect indicator of the structural integrity and aggregation propensity of the βLG variants as described by Keppler et al.^15^. At pH 4.6, which is close to the isoelectric point (pI) of βLG, reduced electrostatic repulsion can promote protein precipitation. Differences in pI among variants may therefore contribute to differences in acid solubility. The ProteinMPNN variants exhibited acid solubility ranging from 90.7 ± 2.7% to 95.6 ± 2.6%, comparable to WT_B (93.1 ± 4.0%). In contrast, the BCAA-stabilized variants showed lower acid solubility, ranging from 67.7 ± 7.2% to 88.6 ± 0.5% (**Fig. 4A**). BCAA variants had a predicted pI only 0.18 pH units above the assay pH, suggesting that their proximity to the pI may have contributed to their reduced solubility. Overall, the high acid solubility observed for the ProteinMPNN variants, even variants with an pI near the test pH, suggests that their structural integrity was retained under the assay conditions, whereas the lower solubility of the BCAA variants may reflect a combination of pI proximity and variant-specific structural or aggregation properties. The predicted pI varied between 4.7 and 5.5 among the variants. The Ip values were computationally predicted and may deviate from experimentally determined values for individual variants.

**Figure 4.**
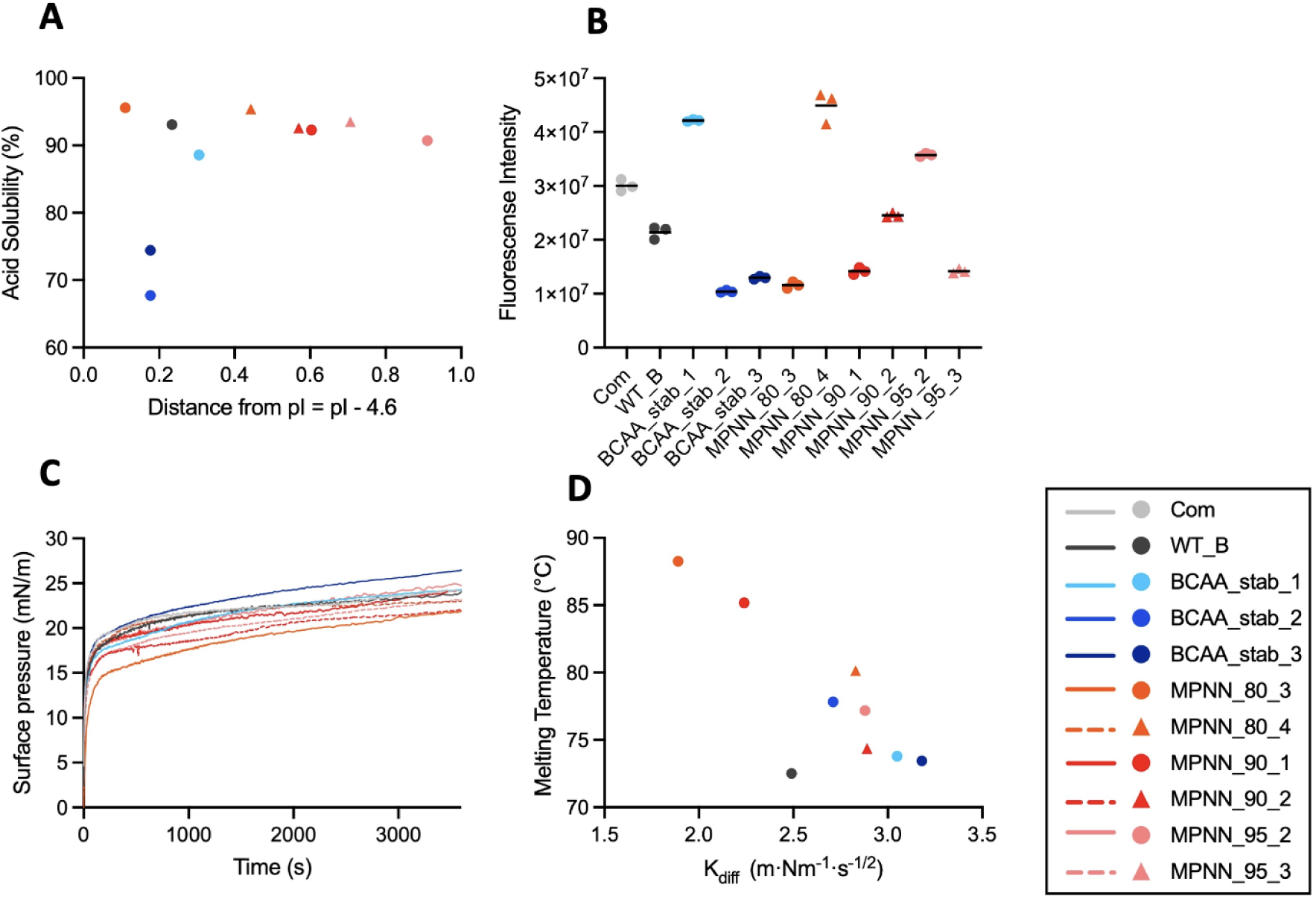
Surface properties, acid solubility, and interfacial behavior of β-lactoglobulin variants. **(A)** Acid solubility as a function of the distance between the experimental pH (pH 4.6) and the predicted isoelectric point (pI) of each β-lactoglobulin variant. **(B)** Surface hydrophobicity of β-lactoglobulin variants assessed by ANS fluorescence intensity. The results are normalized by the protein concentration. **(C)** Surface pressure as a function of time for β-lactoglobulin variants. **(D)** Melting temperature (Tm) as a function of the interfacial adsorption rate constant (Kdiff) (ρ = −0.976, *p* < 0.001).

The relative exposure of hydrophobic regions in the βLG variants was assessed using binding of the fluorescent dye 8-Anilino-1-naphthalenesulfonic acid (ANS), where the ANS emission intensity at 475 nm is used as a measure of ANS-accessible hydrophobic regions. The ANS responses varied substantially among the tested proteins, with the largest difference observed between the two ProteinMPNN 80 variants, MPNN_80_3 and MPNN_80_4 (**Fig. 4B**). The commercial reference and βLG-B showed intermediate ANS responses within this range. Both the ProteinMPNN- and BCAA-stabilized variants exhibited variation in ANS emission, indicating differences in the exposure of ANS-accessible hydrophobic regions despite minor changes in protein sequence. Interestingly, this variation did not fully follow the increasing number of hydrophobic residues in the BCAA variants or the surface hydrophobicity visualized from SASA in PyMOL (**Fig. S4**).

Finally, we assessed air-water interfacial behavior of the βLG variants using droplet tensiometry. Surface-pressure profiles over 60 minutes showed small differences in both adsorption kinetics and final interfacial pressure among the βLG variants (**Fig. 4C, Table S7**). The calculated adsorption rate constant K_diff_ is an indicator of the initial rate of protein accumulation at the interface, whereas the final surface pressure reflects the extent of interfacial pressure development at the end of the measurement. The BCAA-stabilized variants and most ProteinMPNN variants showed K_diff_, and final surface pressure values close to those of βLG-B (K_diff_ = 3.05; final surface pressure = 1.56), suggesting that the introduced substitutions had limited effects on interfacial behavior under the conditions tested. The largest deviation was observed for MPNN_80_3, which exhibited a lower K_diff_ (1.89) and final surface pressure (0.99), indicating slower initial adsorption and lower surface-pressure development at the interface. Standard deviations in K_diff_, were small (0.05–0.26), indicating relatively low variation between replicate measurements. Higher thermal stability was associated with slower interfacial adsorption kinetics (Spearman’s ρ = −0.976, *p* < 0.001) (**Fig. 4D**). Variant MPNN_80_3 was exemplary of this relationship, as it was among the most thermally stable variants and exhibited the lowest K_diff_. This trend spanned all sequence variants, complicating attempts to ascribe interfacial behaviour effect to individual substitutions.

Variant BCAA_stab_3 reached the highest surface pressure after 60 min (26.35 mN/m), compared to βLG-B (24.11 mN/m) and the variant with the lowest end pressure, MPNN_80_3, (21.97 mN/m). The distinct interfacial behaviour of BCAA_stab_3 may arise from its higher hydrophobicity, or from its larger monomeric fraction. Oligomeric state may influence adsorption because smaller protein species can diffuse more rapidly to an interface^33^.

### Correlation between computational metrics and protein biophysical properties

To examine relationships between computational descriptors and experimentally measured properties, Spearman’s rank correlation analysis was performed across the studied variants (**Fig. 5A, Table S8**). Predicted structural deviation from βLG-B, expressed as RMSD, was positively correlated with melting temperature (Tm; ρ = 0.664, *p* = 0.018) and negatively correlated with the relative Ellman’s assay response (ρ = −0.811, *p* = 0.004) (**Fig. 5B**). This correlation likely manifests from a correlation between extent of mutation, and thus stability, and the predicted computational structures deviation from the βLG-B structure. The computational descriptors ‘MPNN score,’ and ‘disorder score,’ showed little or no correlation with thermal stability. RMSD was also positively correlated with protein yield (ρ = 0.720, *p* = 0.004). Interestingly, the computationally calculated fraction of hydrophobic solvent-accessible surface area (hydrophobic SASA ratio), was strongly negatively correlated with protein yield (ρ = −0.850, *p* = 0.001). However, experimentally determined ANS fluorescence was not strongly associated with the predicted hydrophobic SASA ratio.

**Figure 5.**
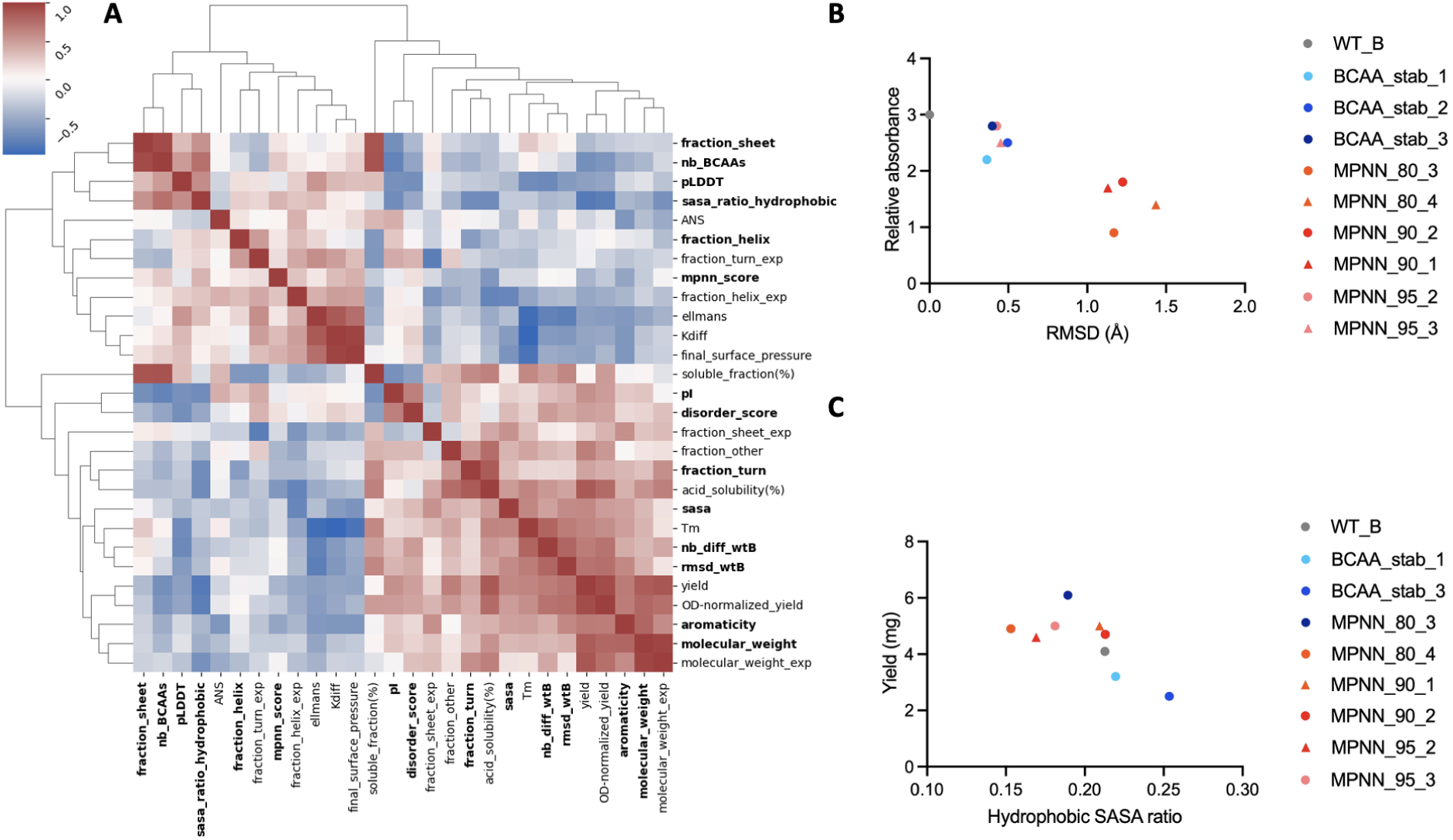
Relationships between computational descriptors and experimentally measured properties of β-lactoglobulin variants. **(A)** Spearman correlation matrix of computational descriptors and experimentally measured properties across β-lactoglobulin variants. **(B)** Relative Ellman’s assay absorbance as a function of predicted structural deviation from WT-B, expressed as root-mean-square deviation (RMSD) (ρ = −0.811, *p* = 0.004). **(C)** Hydrophobic solvent-accessible surface area (SASA) ratio as a function of yield (ρ = −0.850, *p* = 0.001).

## Discussion

In this work we used two different protein design tools to mutate βLG. The objectives of these two tools are different. ProteinMPNN primarily searches for sequences compatible with the desired backbone structure, often resulting in proteins with increased structural stability. Our work shows that ProteinMPNN reliably produces βLG mutants that are more thermally stable (up to 15 °C) and at higher production yields than natural variants. Remarkably, all variants existed as dimers and with similar helical and sheet content, despite up to 20% of the sequence being mutated. Improved thermal stability is relevant for βLG because it is sensitive to heat, with irreversible structural changes and aggregation can occur above approximately 60–70 °C, temperatures that are common in food processing^34^. ProteinMPNN-based redesign has recently improved the thermal stability of several enzymes such as TEV protease and pectinase while maintaining or improving catalytic activity^2,35,36^. Our results show for the first time how ProteinMPNN can be applied to a major functional food protein to improve thermal stability and solubility, while also revealing how these changes relate to other food-relevant functional properties

In the PyRosetta BCAA approach, we specified only BCAA insertion and selected mutants with increased predicted stability, as estimated from simulations of low energy structures. We were able to significantly increase BCAA content from 41 to 49 amino acids while retaining protein thermal stability. Increasing BCAA content is likely destabilizing for protein structure. Despite this, the computational stability predictions were consistent with experimental recovery: variants predicted to be stabilized (Cohen’s *d* = −12.59 to −9.29) could be recovered as soluble protein, whereas the two variants predicted to be destabilized (Cohen’s *d* = 7.9 and 9.7) did not yield detectable soluble protein. This is consistent with previous Rosetta-based protein design studies in which increased predicted or experimentally determined stability has been associated with improved soluble expression^37^. Our work thus provides a proven computational strategy for increasing nutritional content of other food proteins, namely extensive, combinatorial substitution and filtering using physics-based energy models.

Modulation of BCAA content also affected predicted surface hydrophobicity, which influenced protein yield and acid solubility. Whether altered surface hydrophobicity is desirable depends on the application. The reduced yield of BCAA variants may also be the result of host strain limitations. Overexpression of proteins with a high BCAA content has been shown to hinder growth^38^. Further, it is difficult to produce high yields of BCAA due to feedback inhibition and issues balancing growth and BCAA synthesis^39^. βLG already has a high BCAA content and increasing it further could add additional strain on the cells and reduce growth, making it difficult to reach high enough amounts to purify. Dong et al. showed that metabolic engineering of *Pichia pastoris* can increase flux by overexpressing an enzyme in BCAA production pathway^40^.

A key motivation of this work was to extend protein characterisation from thermal stability to other relevant biophysical properties, such as aggregation and interfacial behavior. Protein adsorption at the air–water interface involves diffusion to the interface followed by adsorption and structural rearrangements^41^. Previous studies have shown that βLG is not fully unfolded during adsorption but undergoes limited conformational changes at the air–water interface^42,43^. This may explain why the overall differences in interfacial behaviour remained relatively small. In our study, the negative relationship between melting temperature and the diffusion-controlled adsorption rate suggests that the more thermally stable variants may adsorb more slowly at the air–water interface. However, the differences in adsorption behaviour were relatively small compared with the large differences in thermal stability. More variants and targeted single-mutation studies are needed to determine whether thermal stability and adsorption rate are directly related. More complex emergent properties, such as protein aggregation and gelling propensity, are relevant for protein materials but are difficult to predict from sequence alone^44^. Here, labeled datasets linking sequence to experimental characterisation could allow for the derivation of computational predictors.

Whether the increased thermal stability of the redesigned variants affects other nutritional properties of βLG remains to be investigated. Both ligand binding and digestibility depend on protein structure and conformational flexibility. Keppler et al. showed differences in ligand binding among natural βLG variants and suggested that backbone mobility influences ligand-binding ability^22^. Thus, the increased stability of the ProteinMPNN variants could also affect ligand binding, even if the overall βLG fold is retained. In addition, previous studies using the INFOGEST model have shown that structural modification of βLG by heat treatment increased its gastric digestibility^45^. This therefore raises the hypothesis that the increased structural stability of the ProteinMPNN variants may affect their susceptibility to gastrointestinal proteolysis.

## Conclusion

ProteinMPNN enabled extensive sequence modification, with up to 26 amino acid substitutions introduced while maintaining a βLG-like secondary structure and predominantly dimeric state. Several designed variants showed substantially increased thermal stability (up to 14.5 °C higher than WT_B) and increased solubility despite extensive sequence changes, demonstrating the potential of stability-focused deep learning design for food proteins. The PyRosetta-based approach demonstrated that the nutritional composition of βLG can be directly targeted through protein design. Up to eight additional branched-chain amino acids were successfully introduced while retaining the overall secondary structure. However, increasing the BCAA content reduced protein solubility. Together, these results provide a proof of concept that computational protein engineering can extend beyond conventional enzyme applications to the rational redesign of food proteins. Our results highlight the importance of evaluating not only the properties targeted during protein design, but also potential effects on other food functions. The combination of computational design with experimental characterization provides a framework to in the future understand how sequence-level modifications translate into changes in non-targeted properties, such as ligand binding and gastrointestinal digestibility.

## Material and Methods

### Computational design of βLG variants

The bovine βLG variant B reference sequence was downloaded from UniProt (accession id: P02754)^46^. The signal sequence (residues 1-16) was manually removed before computational analysis and sequence position was renumbered. ProteinMPNN was run using default parameters. In PyRosetta workflows, to account both for structure prediction artifacts and post-mutation steric constraints, all structures were energy-minimized using the FastRelax function^47^ before being scored with Ref2015. Because the FastRelax function produces a stochastic output, we systematically computed 50 relaxation replicates. To enable unbiased comparative analysis of the scoring results, we also replicated relaxation and scoring of the unmutated structure 50 times.

### Computational derivation of biophysical parameters

Unless otherwise specified, physicochemical and sequence-derived properties, including pI, molecular weight, aromaticity, and secondary-structure-related quantities, were calculated using the ProteinAnalysis module implemented in Biopython. SASA was calculated using the Shrake–Rupley rolling-ball algorithm as implemented in Biopython, and the relative contribution of hydrophobic residues was calculated as the ratio of the SASA of alanine, valine, isoleucine, leucine, methionine, phenylalanine, tyrosine, and tryptophan residues to the total SASA. Intrinsic disorder was evaluated using AIUPred, while ProteinMPNN confidence scores were recomputed for all sequences using the model’s score_only functionality. Mean pLDDT values were obtained directly from the corresponding structure-prediction outputs. Given the relatively small number of candidates, structures were subsequently re-predicted using the AlphaFold 3 web-server implementation, allowing structure-based metrics to be recalculated from these higher-quality predictions. This provided a consistent structural basis for the comparative analysis while remaining computationally tractable within the scope of the study.

### Expression of βLG variants

A commercial sample (Com), containing a mixture of isoforms A and B, was used as a positive control for all the methods. The sample was prepared by dissolving lyophilized powder of ßLG from bovine milk (CAS 9045-23-2, Sigma-Aldrich, Missouri, USA). For recombinant variants, DNA fragments encoding the β-lactoglobulin variants (codon optimized for bacterial expression) were purchased from Integrated DNA Technologies (IDT) as gBlocks. The genes encoding the βLG variants included an N-terminal Cytiva Protein Select affinity tag and were synthesized with overhangs compatible with Gibson Assembly and cloned into the pET-28a expression vector. The Gibson Assembly products were transformed into *Escherichia coli* XL1-Blue competent cells by heat-shock transformation and plated on LB agar containing the appropriate antibiotic. Positive colonies were verified by Sanger sequencing (Eurofins Genomics). Plasmids were isolated using the GeneJET Plasmid Miniprep Kit (Thermo Fisher Scientific, Vilnius, Lithuania) and transformed into *E. coli* SHuffle T7 competent cells. Single colonies were used to prepare glycerol stocks (25% glycerol) and overnight starter cultures for protein expression. Protein expression was carried out in LB medium inoculated with 1% (v/v) overnight culture and incubated at 30 °C with shaking until an OD₆₀₀ of 0.4–0.8 was reached. Protein expression was induced with 0.2 or 0.4 mM IPTG, followed by incubation at 16 °C for 16–24 h with shaking. Cells were harvested by centrifugation (4500 × *g*, 15 min, 4 °C) and stored at −20 °C until purification. Cell pellets obtained from 0.25, 0.50, or 1.0 L cultures were resuspended in 20 mM sodium phosphate buffer containing 200 mM NaCl (pH 7.4) and disrupted by sonication (10 s on/10 s off, 180 cycles). The resuspension volume was adjusted according to culture size (20 mL for pellets from 0.25 L cultures and 40 mL for pellets from 1.0 L cultures). B-PER Complete Bacterial Protein Extraction Reagent (Thermo Fisher Scientific, Illinois, USA) was added at 20 mL per liter of original culture volume, and samples were incubated for 30 min at room temperature. Cell debris was removed by centrifugation (14,000 × *g*, 15 min, 4 °C), and the soluble fraction was filtered through a 0.2 μm membrane filter.

### Protein purification

Proteins were purified using an ÄKTA start chromatography system (Cytiva, Uppsala, Sweden) equipped with a 1 mL HiTrap Protein Select column. The column was equilibrated, washed, and eluted using a 20 mM sodium phosphate buffer containing 200 mM NaCl (pH 7.4). A static binding step of 3.5 h was introduced between the wash and elution steps. Expression, solubility, and purification were monitored by reducing SDS-PAGE using 50 mM dithiothreitol (DTT). Soluble fractions, insoluble fractions, and purified protein fractions were analyzed after each purification to assess protein expression and purity.

### Protein concentration determination

Protein concentrations were determined using a NanoPhotometer NP80 (Implen, Munich, Germany). Concentrations were calculated using an extinction coefficient of 0.937 L g⁻¹ cm⁻¹ for all the other variants except 1.019 L g⁻¹ cm⁻¹ and 1.307 L g⁻¹ cm⁻¹ for variants MPNN_90_1 and MPNN_80_3 respectively, for which variant-specific extinction coefficients predicted by ProtParam were used.

### Secondary structure by Circular dichroism

Circular dichroism (CD) spectroscopy was used to assess the secondary structure of purified proteins. Prior to analysis, proteins were buffer-exchanged into 10 mM sodium phosphate buffer (pH 8.0) using 10 kDa molecular weight cutoff centrifugal filters and at least seven buffer exchange volumes. Samples were diluted to 0.2 mg mL⁻¹ in Milli-Q water immediately before analysis. CD spectra were recorded on a Chirascan spectropolarimeter (Applied Photophysics, Surrey, UK) using a 1 mm pathlength cuvette at 20 °C over a wavelength range of 180–260 nm with a 1 nm step size and 1 s integration time. Three technical replicates were recorded for each sample. Spectra were averaged and converted from ellipticity (mdeg) to mean residue ellipticity assuming a mean residue mass of 113 Da (18.3 kDa/162 residues). Secondary structure content was estimated using the BeStSel web server^48^.

### MALDI-TOF Mass Spectrometry

Matrix-assisted laser desorption/ionization time-of-flight mass spectrometry (MALDI-TOF MS) was used to determine the molecular masses of purified proteins. The buffer-exchanged samples prepared for CD analysis were analyzed using a 4800 MALDI TOF/TOF Analyzer (Applied Biosystems, Massachusetts, USA). Spectra were acquired over an *m/z* range of 5,000–20,000. Measured molecular masses were compared with theoretical masses calculated using ProtParam^49^.

### Size Exclusion Chromatography

High-performance size exclusion chromatography (HPLC-SEC) was performed to assess protein purity and oligomeric state. Samples were diluted to approximately 1 mg mL⁻¹ in 20 mM sodium phosphate buffer containing 200 mM NaCl (pH 7.4) and analyzed using an Agilent 1100 HPLC system (Agilent Technologies, California, USA) equipped with TSKgel G3000SWXL (5 μm, 300 × 7.8 mm) and TSKgel G2000SWXL (5 μm, 300 × 7.8 mm) columns. The mobile phase consisted of 30% acetonitrile containing 0.1% trifluoroacetic acid, and separations were performed at a flow rate of 1.5 mL min⁻¹ with the column maintained at 30 °C. Elution was monitored by absorbance at 214 nm. Species with apparent molecular masses greater than 60 kDa eluted in the high-molecular-weight region.

### Nano-Differential Scanning Fluorimetry

NanoDSF was used to assess the thermal stability of purified proteins. Samples were diluted to approximately 0.5 mg mL⁻¹ in 10 mM sodium phosphate buffer containing 100 mM NaCl (pH 7.4). Three technical replicates (approximately 20 μL each) were loaded into capillaries and analyzed using a Prometheus NT.48 instrument (NanoTemper Technologies, Munich, Germany). Fluorescence at 330 and 350 nm was monitored while the temperature was increased from 20 to 95 °C at a rate of 1.0 °C min⁻¹. The onset temperature (Tₒ) and melting temperature (T◻) were determined from the F350/F330 fluorescence ratio using PR.ThermControl software (NanoTemper Technologies).

### Ellman’s Assay

Free thiol groups were quantified using Ellman’s assay^50^. A 10 mM solution of 5,5′-dithiobis-(2-nitrobenzoic acid) (DTNB; Ellman’s reagent) was freshly prepared in a 100 mM sodium phosphate buffer (pH 7.4) on the day of the experiment. Protein concentrations were determined prior to analysis. Protein samples in a 20 mM sodium phosphate buffer containing 200 mM NaCl (pH 7.4) were analyzed either without heat treatment or after incubation at 63 or 72 °C for 15 min, 30 min, or 1 h. Immediately after heating, samples were cooled on ice. For each measurement, 15 μL of protein sample was mixed with 85 μL of 10 mM sodium phosphate buffer containing 100 mM NaCl (pH 7.4) and 2 μL of 10 mM DTNB in a 96-well microplate. Absorbance at 412 nm was recorded for 6 min using a spectrophotometer (SpectraMaxR i3x, Molecular Devices or BioTek Epoch 2 microplate reader, Agilent) Buffer containing DTNB without protein served as the blank. All measurements were performed in technical triplicate. The average absorbance over the first 4 min was used for analysis. Free thiol concentrations were calculated using an optical pathlength of 0.276 cm, a molar extinction coefficient of 13,600 M⁻¹ cm⁻¹ for TNB at 412 nm, and a molecular weight of 18.3 kDa for β-lactoglobulin.

### Acid solubility

Acid solubility was determined using a modified version of the method described by Keppler et al.^15^ and Sava et al.^51^. Protein samples were concentrated to approximately 3–5 mg mL⁻¹ and diluted 10-fold into 10 mM sodium acetate containing 50 mM NaCl (pH 4.6). After incubation at 20 °C with shaking (100 rpm) for 60 min, samples were centrifuged at 20,800 × *g* for 60 min. Protein concentrations in the supernatant were determined using a NanoPhotometer NP80 (Implen, Munich, Germany), and acid solubility was calculated as the percentage of soluble protein relative to the initial protein concentration.

### Surface hydrophobicity by ANS

Surface hydrophobicity was determined using 8-anilino-1-naphthalenesulfonic acid (ANS) fluorescence as described in Deng et al.^34^. A 4 mM ANS working solution was prepared from an 8 mM stock solution in a 100 mM sodium phosphate buffer (pH 7.4). Protein samples were diluted to 0.1–0.2 mg mL⁻¹ in 10 mM sodium phosphate buffer containing 100 mM NaCl (pH 7.4). ANS solution (10 μL) was added to 1 mL of protein sample, followed by incubation in the dark for 15 min at room temperature. Aliquots (200 μL) were transferred in technical triplicate to a black 96-well microplate. Fluorescence was measured using a plate reader with excitation at 390 nm and emission recorded from 460 to 500 nm at 10 nm intervals. A spectrophotometer (SpectraMaxR i3x, Molecular Devices or BioTek Epoch 2 microplate reader, Agilent) with excitation and emission bandwidths of 9 and 15 nm was used, respectively.

### Droplet Tensiometry

Dynamic surface tension measurements were performed using a TRACKER drop tensiometer (Teclis Scientific, Civrieux d’Azergues, France) employing the pendant drop method. A 6 μL droplet of protein solution (0.1 mg mL⁻¹ in 20 mM sodium phosphate buffer containing 200 mM NaCl, pH 7.4) was monitored at room temperature for 1 h. All samples were measured in duplicate, except variants MPNN_90_2 and MPNN_95_3, which were measured overnight. Surface tension was recorded once per second for the first 1000 s and every 10 s thereafter. Between measurements, the syringe and needle were cleaned with ethanol and Milli-Q water. Buffer measurements were performed to verify the absence of surface-active contaminants. Surface pressure was calculated as the difference between the surface tension of the buffer and that of the protein solution. Duplicate measurements were interpolated to a common time axis before averaging. Curves were smoothed using a 7-point moving average for visualization. The initial adsorption rate (*K*diff) was determined using the Ward–Tordai approach as described by Kontogiorgos et al.^52^. Surface pressure (π) was plotted against the square root of time (*t*1/2), and *K*diff was calculated as the slope of the initial linear region.

## Acknowledgements

We are grateful to Olivia Liljefors (KTH) for assistance in purifying recombinant βLG. We are also grateful to Boxin Deng and Vega Mortes for their assistance with the drop tensiometer measurements in Wageningen.

## Supplementary Tables and Figures

**Table S1.**
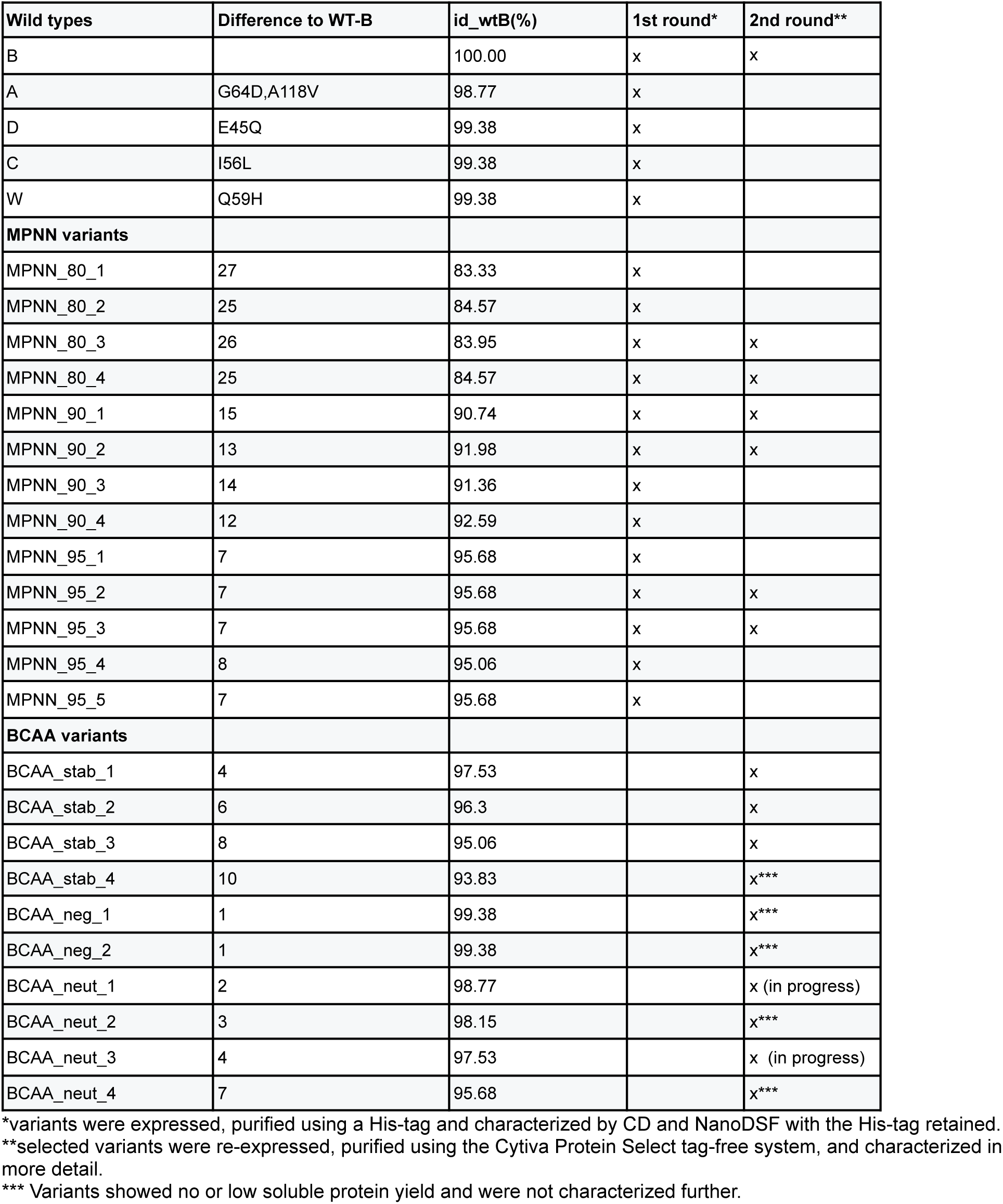
β-Lactoglobulin variants generated and evaluated in this study. Variants are shown with their number of amino acid substitutions and sequence identity relative to wild-type βLG B (WT-B).

**Table S2.** Computational structural and physicochemical parameters of the β-lactoglobulin (βLG) variants.

| βLG Variant | Fraction<br>_helix, % | Fraction_<br>turn, % | Fraction<br>_sheet,<br>% | Isoelectric<br>point, pI | Molecular<br>weight, Da | Aromaticity, | SASA | SASA_ratio_<br>hydrophobic,<br>Å<sup>2</sup> |
| --- | --- | --- | --- | --- | --- | --- | --- | --- |
| WT-B | 0.444 | 0.21 | 0.364 | 4.834 | 18281.005 | 0.062 | 8401.36 | 0.213 |
| A | 0.438 | 0.21 | 0.37 | 4.76 | 18367.095 | 0.062 | 8382.86 | 0.203 |
| C | 0.451 | 0.21 | 0.364 | 4.834 | 18281.005 | 0.062 | 8328.33 | 0.218 |
| D | 0.438 | 0.21 | 0.364 | 4.905 | 18280.021 | 0.062 | 8163.55 | 0.21 |
| W | 0.444 | 0.21 | 0.364 | 4.924 | 18290.016 | 0.062 | 8249.34 | 0.205 |
| MPNN_80_1 | 0.407 | 0.222 | 0.383 | 5.067 | 18592.362 | 0.08 | 8577.79 | 0.189 |
| MPNN_80_2 | 0.395 | 0.222 | 0.414 | 4.9 | 18316.109 | 0.068 | 8401.7 | 0.175 |
| MPNN_80_3 | 0.395 | 0.228 | 0.389 | 4.71 | 18513.176 | 0.074 | 8572.6 | 0.189 |
| MPNN_80_4 | 0.414 | 0.228 | 0.389 | 5.043 | 18325.078 | 0.062 | 8384.19 | 0.153 |
| MPNN_90_1 | 0.451 | 0.204 | 0.352 | 5.203 | 18359.164 | 0.068 | 8389.58 | 0.209 |
| MPNN_90_2 | 0.444 | 0.204 | 0.364 | 5.169 | 18219.07 | 0.062 | 8429.87 | 0.213 |
| MPNN_90_3 | 0.444 | 0.204 | 0.358 | 5.067 | 18372.163 | 0.068 | 8394.81 | 0.189 |
| MPNN_90_4 | 0.444 | 0.216 | 0.352 | 5.572 | 18252.1 | 0.062 | 8474.68 | 0.182 |
| MPNN_95_1 | 0.444 | 0.21 | 0.352 | 5.306 | 18254.975 | 0.062 | 8342.85 | 0.176 |
| MPNN_95_2 | 0.444 | 0.21 | 0.346 | 5.511 | 18310.057 | 0.062 | 8226.76 | 0.169 |
| MPNN_95_3 | 0.432 | 0.216 | 0.352 | 5.306 | 18324.04 | 0.062 | 8526.36 | 0.181 |
| MPNN_95_4 | 0.438 | 0.216 | 0.352 | 5.306 | 18312.026 | 0.062 | 8396.98 | 0.175 |
| MPNN_95_5 | 0.438 | 0.21 | 0.352 | 5.306 | 18281.015 | 0.062 | 8411.39 | 0.187 |
| BCAA_stab_1 | 0.444 | 0.191 | 0.389 | 4.905 | 18291.172 | 0.062 | 8228.95 | 0.22 |
| BCAA_stab_2 | 0.432 | 0.204 | 0.401 | 4.777 | 18259.17 | 0.062 | 8469.67 | 0.252 |
| BCAA_stab_3 | 0.444 | 0.198 | 0.407 | 4.777 | 18297.304 | 0.062 | 8174.37 | 0.253 |
| BCAA_stab_4 | 0.444 | 0.191 | 0.414 | 4.777 | 18311.374 | 0.062 | 8416 | 0.252 |
| BCAA_neg_1 | 0.444 | 0.204 | 0.37 | 4.834 | 18280.06 | 0.062 | 8427.5 | 0.237 |
| BCAA_neg_2 | 0.444 | 0.204 | 0.37 | 4.834 | 18337.112 | 0.062 | 8374.04 | 0.213 |
| BCAA_neut_1 | 0.438 | 0.21 | 0.377 | 4.746 | 18236.993 | 0.062 | 8140.7 | 0.231 |
| BCAA_neut_2 | 0.438 | 0.21 | 0.377 | 4.664 | 18249.003 | 0.062 | 8325.86 | 0.247 |
| BCAA_neut_3 | 0.438 | 0.21 | 0.383 | 4.806 | 18187.02 | 0.056 | 8263.26 | 0.242 |
| BCAA_neut_4 | 0.438 | 0.198 | 0.395 | 4.792 | 18195.128 | 0.056 | 8197.23 | 0.266 |

**Table S3.** Experimental molecular weight, protein yield, and solubility of β-lactoglobulin (βLG) variants.

| **βLG variants** | **Measured molecular weight (Da)** | **Difference to computed molecular weight (Da)** | **βLG yield\* (mg)** | **Soluble fraction\*\* (%)** |
| --- | --- | --- | --- | --- |
| Com | 18327 | n.a. | n.a. | n.a. |
| WT-B | 18260 | 22 | 4.1 | 72 |
| BCAA_stab_1 | 18245 | 47 | 3.2 | 59 |
| BCAA_stab_2 | 17504 | 755 |  | 64 |
| BCAA_stab_3 | 18267 | 31 | 2.5 | 71 |
| MPNN_80_3 | 18472 | 42 | 6.1 | 91 |
| MPNN_80_4 | 18301 | 25 | 4.9 | 93 |
| MPNN_90_1 | 18341 | 18 | 5.0 | 58 |
| MPNN_90_2 | 18184 | 35 | 4.7 | 72 |
| MPNN_95_2 | 18287 | 24 | 4.6 | 71 |
| MPNN_95_3 | 18310 | 15 | 5.0 | 68 |

\* βLG yield represents the amount of purified βLG recovered from 250 mL of expression culture, based on absorbance at 280 and purity estimated by SEC peak area (based on one replicate).
\*\* Soluble fraction represents the percentage of βLG in the soluble fraction after lysis (based on one replicate).

**Table S4.** Areas of the peaks in HPLC-SEC measurements. Monomer and dimer percentages are calculated based on peak areas, excluding the peaks over 60 kDa. n.a. signifies that a peak was not detected.

| **βLG variants** | **>60 kDa**<br>**(mAU·min)** | **Tagged**<br>**βLG**<br>**(mAU·min)** | **Dimer**<br>**(mAU·min)** | **Monomer**<br>**(mAU·min)** | **Dimer**<br>**(%)** | **Monomer**<br>**(%)** | **Purity\* (%)** |
| --- | --- | --- | --- | --- | --- | --- | --- |
| Com | 3.09 | n.a. | 78.4 | n.a. | 100 | 0 | 96 |
| WT_A | 3.5 | n.a. | 19.15 | 0.34 | 98 | 2 | 85 |
| WT_B | 2.74 | n.a. | 50.83 | 1.39 | 97 | 3 | 95 |
| WT_C | 11.14 | n.a. | 9.55 | 1.05 | 90 | 10 | 49 |
| WT_W | 3.55 | n.a. | 82.52 | 2.31 | 97 | 3 | 96 |
| BCAA_stab_1 | 7.57 | n.a. | 49.86 | 1.39 | 97 | 3 | 87 |
| BCAA_stab_3 | 9.54 | n.a. | 21.77 | 2.49 | 90 | 10 | 72 |
| MPNN_80_3 | 1.63 | 11.85 | 40.46 | 2.25 | 96 | 4 | 97 |
| MPNN_80_4 | 2.56 | 4.15 | 57.94 | 2.56 | 96 | 4 | 96 |
| MPNN_90_1 | 1.63 | n.a. | 46.51 | 2 | 96 | 4 | 97 |
| MPNN_90_2 | 1.76 | n.a. | 59.7 | 1.81 | 97 | 3 | 97 |
| MPNN_95_2 | 2.71 | n.a. | 45.38 | 2.08 | 96 | 4 | 95 |
| MPNN_95_3 | 2.1 | n.a. | 52.98 | 2.03 | 96 | 4 | 96 |

\* Tagged βLG has not been regarded as an impurity.

**Table S5.**
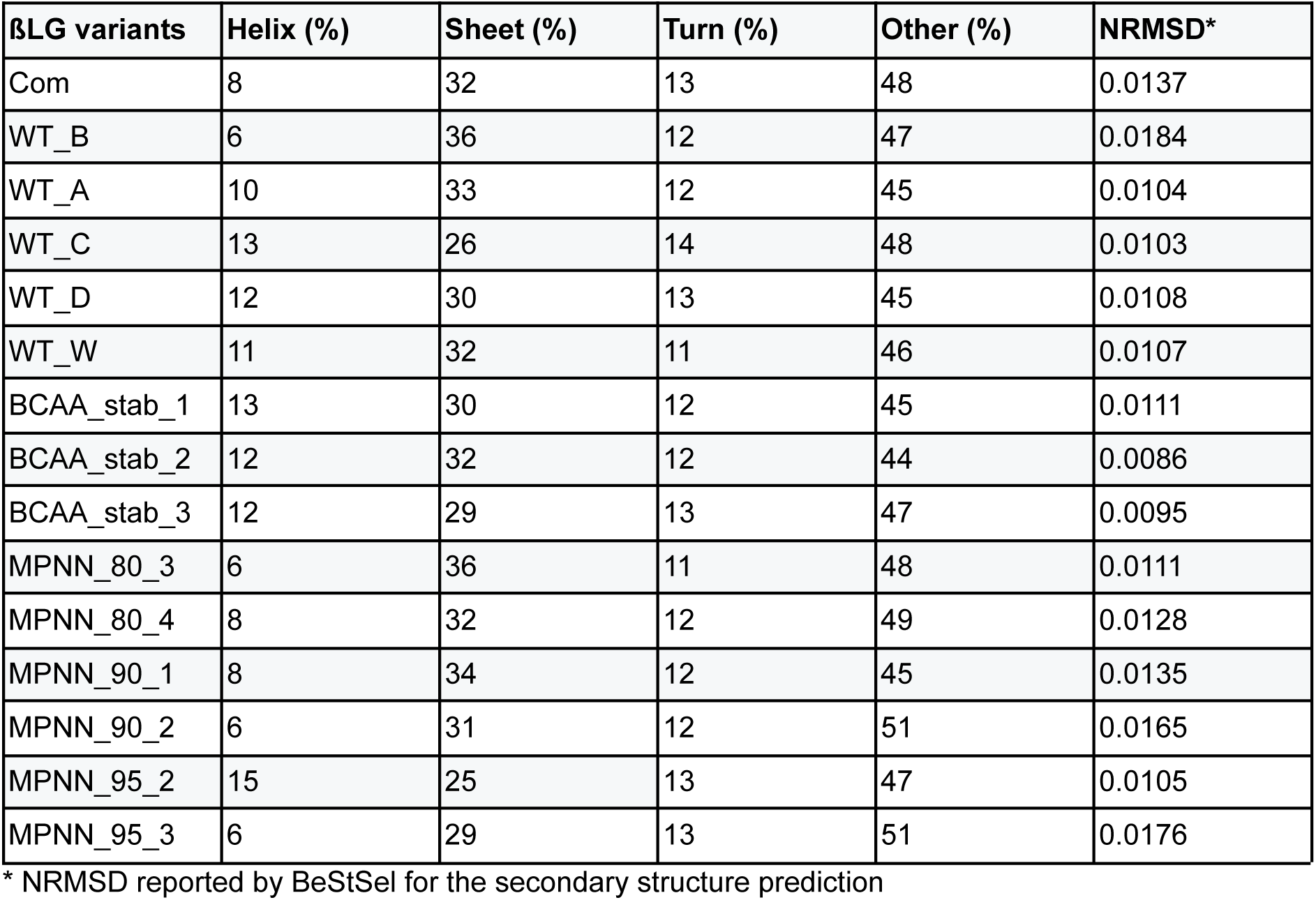
The proportions of α-helix, β-sheet, turn, and other secondary structure elements were estimated from the CD spectra using BeStSel. *NRMSD, normalized root-mean-square deviation, indicates the goodness of fit between the experimental and fitted spectra.

**Table S6.** Thermal stability, thiol accessibility, acid solubility, and surface hydrophobicity of β-lactoglobulin (βLG) variants.

| **βLG variants** | **Melting temperature (°C)** | **Ellman's Fluorecense\*** | **Acid solubility (%)** | **ANS fluorescence** |
| --- | --- | --- | --- | --- |
| Com | 72.5 ±0.1 | 3.6 ±0.34 | 97.8 ±2.8 | 30057810 ±1096723 |
| WT_B | 73.8 ±2.0 | 3.0 ±0.22 | 93.1 ±4.0 | 21414974 ±1190387 |
| WT_A | 72.4 ±0.1 | | | |
| WT_C | 71.0 ±2.5 | | | |
| WT_D | 71.8 ±0.3 | | | |
| WT_W | 72.1±0.0 | | | |
| BCAA_stab_1 | 77.8 ±0.2 | 2.2 ±0.11 | 88.6 ±0.5 | 42162901 ±175770 |
| BCAA_stab_2 | | 2.5 ±0.13 | 67.7 ±7.2 | 10394158 ±231957 |
| BCAA_stab_3 | 73.5 ±0.1 | 2.8 ±0.19 | 74.4 ±2.0 | 12959586 ±272104 |
| MPNN_80_3 | 88.3 ±0.3 | 0.9 ±0.11 | 95.6 ±2.6 | 11593142 ±637600 |
| MPNN_80_4 | 80.1 ±0.1 | 1.4 ±0.14 | 95.4 ±0.8 | 44918898 ±2953860 |
| MPNN_90_1 | | 1.7 ±0.31 | 92.3 ±2.0 | 14192610 ±636737 |
| MPNN_90_2 | 85.2 ±0.2 | 1.8 ±0.28 | 92.6 ±3.3 | 24547403 ±432279 |
| MPNN_95_2 | 74.4 ±1.3 | 2.8 ±0.15 | 90.7 ±2.7 | 35747147 ±308845 |
| MPNN_95_3 | 77.2 ±0.1 | 2.5 ±0.23 | 93.5 ±2.8 | 14204755 ±395147 |

\*Relative to unheated sample after heating 72°C, 30 min

**Table S7.** Interfacial adsorption properties of β-lactoglobulin (βLG) variants at the air–water interface. The diffusion-controlled adsorption rate constant (Kdiff) and final surface pressure were determined from dynamic surface tension measurements at 60 min.

| **βLG variants** | **$K_{diff}$**<br>**($\text{mN}\cdot\text{m}^{-1}\cdot\text{s}^{-1/2}$)** | **Final Surface Pressure**<br>**($\text{mN/m}$)** |
| --- | --- | --- |
| Com | $2.49 \pm 0.21$ | $24.2 \pm 1.11$ |
| WT_B | $3.05 \pm 0.08$ | $24.1 \pm 0.22$ |
| BCAA_stab_1 | $2.71 \pm 0.26$ | $24.3 \pm 0.06$ |
| BCAA_stab_3 | $3.18 \pm 0.13$ | $26.4 \pm 0.14$ |
| MPNN_80_3 | $1.89 \pm 0.08$ | $22.0 \pm 0.28$ |
| MPNN_80_4 | $2.83 \pm 0.2$ | $23.2 \pm 0.05$ |
| MPNN_90_1 | $2.69 \pm 0.23$ | $24.2 \pm 0.12$ |
| MPNN_90_2 | $2.24 \pm 0.05$ | $22.3 \pm 0.27$ |
| MPNN_95_2 | $2.89 \pm 0.14$ | $24.7 \pm 0.04$ |
| MPNN_95_3 | $2.88 \pm 0.14$ | $23.2 \pm 0.50$ |

**Table S8.** *p*-values from Spearman’s rank correlation analysis. The table reports the *p*-values corresponding to the pairwise Spearman correlations visualized in the correlation heatmap in Figure 5.

|  | nb_diff_wtB | SASA | rmsd_wtB | fraction_helix | fraction_turn | fraction_sheet | pl | Molecular_weight | aromaticity |
| --- | --- | --- | --- | --- | --- | --- | --- | --- | --- |
| nb_diff_wtB | 0 |  |  |  |  |  |  |  |  |
| SASA | 0.011 | 0 |  |  |  |  |  |  |  |
| rmsd_wtB | 0 | 0.001 | 0 |  |  |  |  |  |  |
| fraction_helix | 0.188 | 0.166 | 0.338 | 0 |  |  |  |  |  |
| fraction_turn | 0.296 | 0.092 | 0.305 | 0 | 0 |  |  |  |  |
| fraction_sheet | 0.453 | 0.79 | 0.804 | 0.095 | 0.081 | 0 |  |  |  |
| pl | 0.077 | 0.242 | 0.05 | 0.128 | 0.25 | 0 | 0 |  |  |
| molecular_weight | 0.05 | 0.071 | 0.043 | 0.455 | 0.095 | 0.206 | 0.109 | 0 |  |
| aromaticity | 0.021 | 0.012 | 0.001 | 0.546 | 0.039 | 0.155 | 0.143 | 0 | 0 |
| nb_BCAAs | 0.497 | 0.22 | 0.247 | 0.347 | 0.008 | 0 | 0 | 0.018 | 0.004 |
| pLDDT | 0 | 0.034 | 0.001 | 0.404 | 0.116 | 0.086 | 0 | 0.44 | 0.277 |
| sasa_ratio_hydrophobic | 0.03 | 0.133 | 0.05 | 0.212 | 0 | 0.003 | 0 | 0.011 | 0.027 |
| disorder_score | 0.006 | 0.078 | 0.024 | 0.915 | 0.437 | 0.03 | 0 | 0.269 | 0.327 |
| mpnn_score | 0.978 | 0.28 | 0.855 | 0.518 | 0.031 | 0.516 | 0.602 | 0.142 | 0.003 |
| molecular_weight_exp | 0.143 | 0.629 | 0.668 | 0.2 | 0.04 | 0.402 | 0.893 | 0 | 0.039 |
| yield | 0.009 | 0.023 | 0.004 | 0.815 | 0.098 | 0.165 | 0.039 | 0 | 0.009 |
| soluble_fraction(%) | 0.268 | 0.819 | 0.148 | 0.129 | 0.166 | 0.008 | 0.094 | 0.939 | 0.924 |
| acid_solubility(%) | 0.126 | 0.138 | 0.187 | 0.214 | 0.002 | 0.316 | 0.841 | 0.098 | 0.266 |
| ANS | 0.444 | 0.356 | 0.865 | 0.727 | 0.914 | 0.966 | 0.286 | 0.332 | 0.064 |
| ellmans | 0.007 | 0.29 | 0.004 | 0.571 | 0.368 | 0.781 | 0.775 | 0.115 | 0.053 |
| Tm | 0.002 | 0.036 | 0.018 | 0.314 | 0.521 | 0.405 | 0.555 | 0.24 | 0.114 |
| Kdiff | 0.081 | 0.088 | 0.067 | 0.862 | 0.664 | 0.966 | 0.898 | 0.381 | 0.064 |
| final_surface_pressure | 0.17 | 0.057 | 0.145 | 0.776 | 0.761 | 0.688 | 0.932 | 0.431 | 0.062 |
| fraction_helix_exp | 0.147 | 0.002 | 0.175 | 0.256 | 0.151 | 0.97 | 0.759 | 0.396 | 0.133 |
| fraction_sheet_exp | 0.718 | 0.043 | 0.794 | 0.368 | 0.544 | 0.763 | 0.173 | 0.314 | 0.036 |
| fraction_turn_exp | 0.55 | 0.21 | 0.542 | 0.068 | 0.428 | 0.171 | 0.138 | 0.334 | 0.251 |
| fraction_other | 0.203 | 0.19 | 0.283 | 0.847 | 0.048 | 0.425 | 0.218 | 0.663 | 0.985 |
|  | nb_BCAAs | pLDDT | sasa_ratio_hydrophobic | disorder_score | mpnn_score | molecular_weight_exp | yield | soluble_fraction(%) | acid_solubility(%) |
| nb_BCAAs | 0 |  |  |  |  |  |  |  |  |
| pLDDT | 0.006 | 0 |  |  |  |  |  |  |  |
| sasa_ratio_hydrophobic | 0 | 0.001 | 0 |  |  |  |  |  |  |
| disorder_score | 0.003 | 0 | 0.005 | 0 |  |  |  |  |  |
| mpnn_score | 0.178 | 0.525 | 0.05 | 0.428 | 0 |  |  |  |  |
| molecular_weight_exp | 0.482 | 0.255 | 0.011 | 0.306 | 0.887 | 0 |  |  |  |
| yield | 0.006 | 0.032 | 0 | 0.081 | 0.125 | 0.001 | 0 |  |  |
| soluble_fraction(%) | 0.007 | 0.482 | 0.702 | 0.18 | 0.535 | 0.76 | 0.939 | 0 |  |
| acid_solubility(%) | 0.307 | 0.365 | 0.019 | 0.751 | 0.048 | 0.043 | 0.002 | 0.119 | 0 |
| ANS | 0.932 | 0.546 | 0.46 | 0.732 | 0.898 | 0.17 | 0.544 | 0.432 | 0.831 |
| ellmans | 0.913 | 0.11 | 0.362 | 0.75 | 0.737 | 0.197 | 0.082 | 0.294 | 0.141 |
| Tm | 0.859 | 0.009 | 0.342 | 0.471 | 0.513 | 0.689 | 0.102 | 0.208 | 0.102 |
| Kdiff | 0.73 | 0.356 | 0.798 | 0.546 | 0.381 | 0.516 | 0.137 | 0.482 | 0.244 |
| final_surface_pressure | 0.54 | 0.417 | 0.881 | 0.667 | 0.295 | 0.5 | 0.125 | 0.878 | 0.318 |
| fraction_helix_exp | 0.698 | 0.409 | 0.318 | 0.84 | 0.303 | 0.231 | 0.121 | 0.253 | 0.009 |
| fraction_sheet_exp | 0.67 | 0.737 | 0.648 | 0.342 | 0.483 | 0.255 | 0.51 | 0.482 | 0.214 |
| fraction_turn_exp | 0.119 | 0.615 | 0.497 | 0.187 | 0.562 | 0.464 | 0.841 | 0.119 | 0.364 |
| fraction_other | 0.149 | 0.417 | 0.115 | 0.226 | 0.134 | 0.621 | 0.038 | 0.403 | 0.013 |
|  | ANS | Ellmans | Tm | Kdiff | final_surface_pressure | fraction_helix_exp | fraction_sheet_exp | fraction_turn_exp | fraction_other |
| ANS | 0 |  |  |  |  |  |  |  |  |
| Ellmans | 0.881 | 0 |  |  |  |  |  |  |  |
| Tm | 0.823 | 0.001 | 0 |  |  |  |  |  |  |
| Kdiff | 0.831 | 0.003 | 0 | 0 |  |  |  |  |  |
| final_surface_pressure | 0.635 | 0.022 | 0.002 | 0 | 0 |  |  |  |  |
| fraction_helix_exp | 0.308 | 0.265 | 0.071 | 0.187 | 0.203 | 0 |  |  |  |
| fraction_sheet_exp | 0.488 | 0.204 | 0.297 | 0.139 | 0.232 | 0.012 | 0 |  |  |
| fraction_turn_exp | 0.983 | 0.082 | 0.23 | 0.185 | 0.389 | 0.123 | 0 | 0 |  |
| fraction_other | 0.864 | 0.513 | 0.147 | 0.604 | 0.587 | 0.062 | 0.473 | 0.379 | 0 |

**Figure S1.**
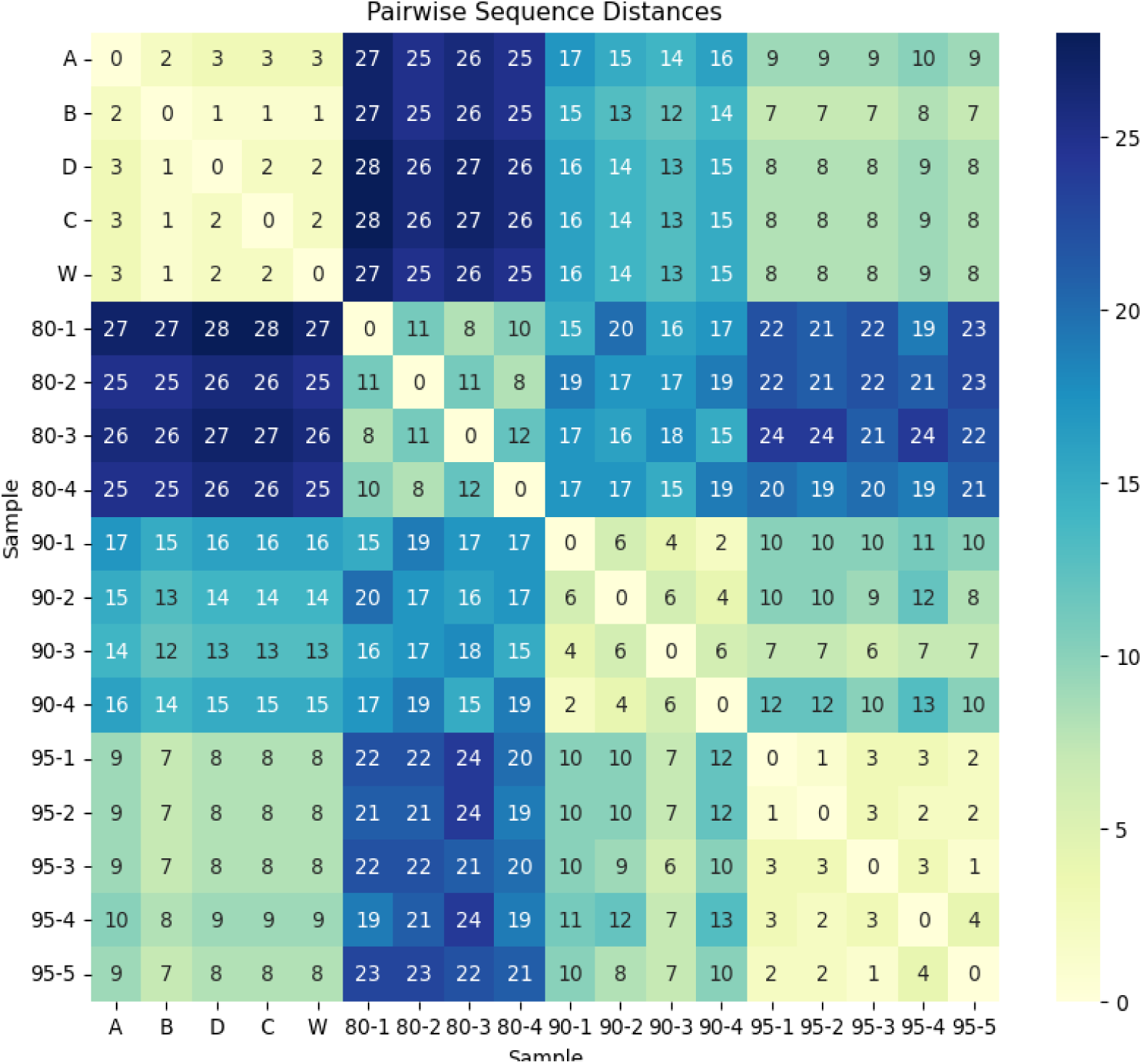
Pairwise sequence distances among wild-type β-lactoglobulin (βLG) sequences and ProteinMPNN-designed variants.

**Figure S2.**
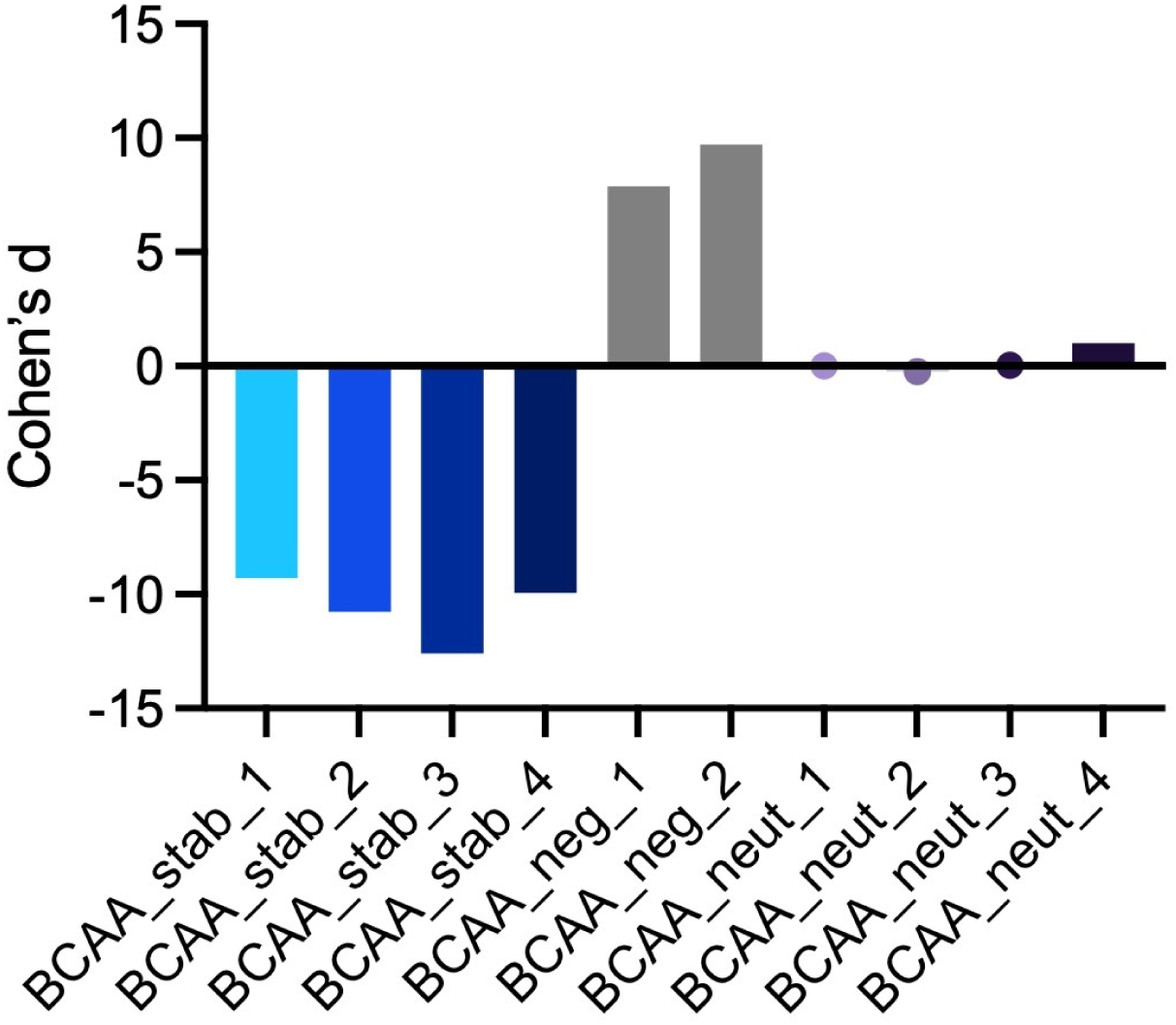
Cohen’s *d* values for the designed BCAA variants relative to wild-type βLG B. Negative values indicate a predicted stabilizing effect, whereas positive values indicate a predicted destabilizing effect.

**Figure S3.**
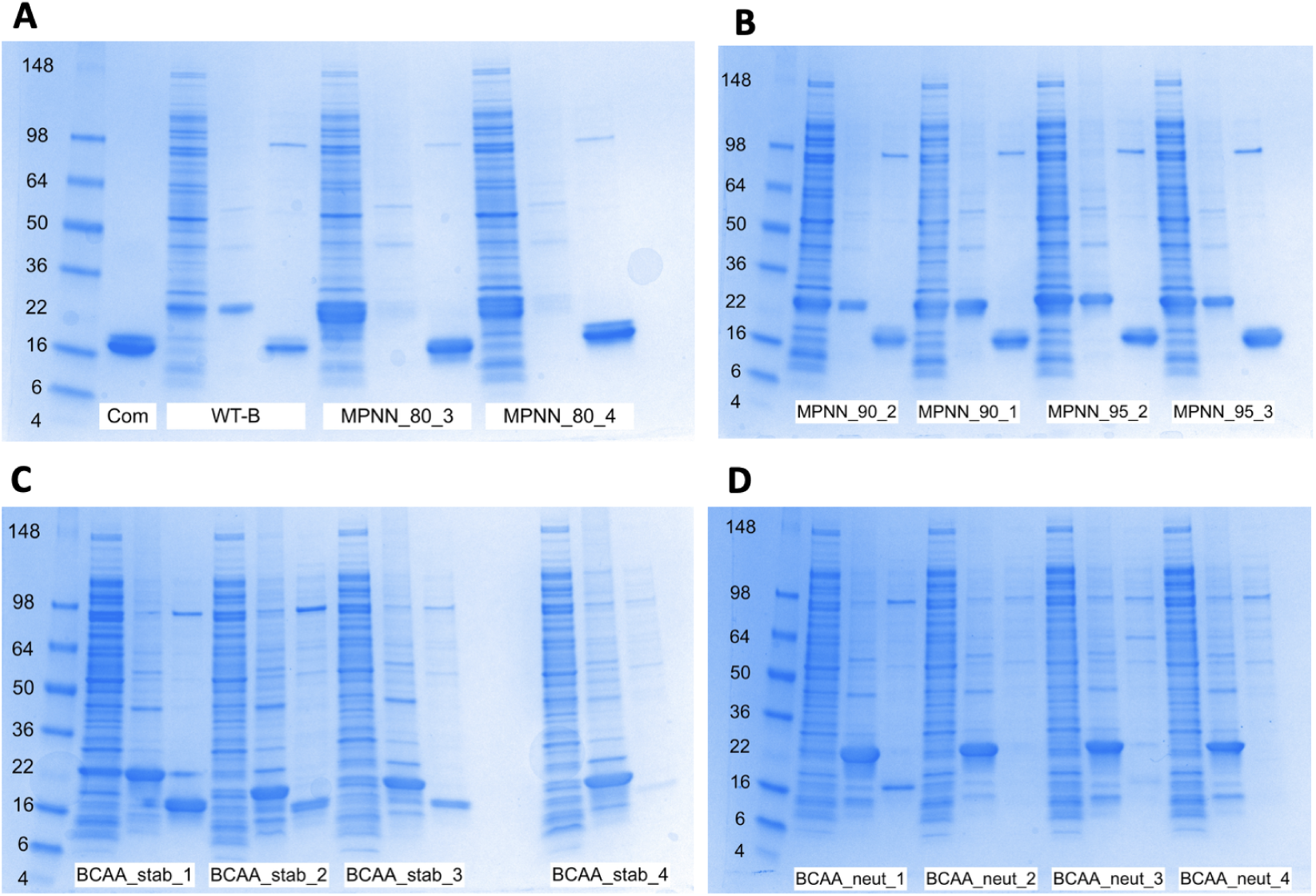
SDS-PAGE analysis of commercial β-lactoglobulin (βLG), wild-type βLG-B (WT-B), ProteinMPNN-designed variants, and BCAA-enriched variants. For WT-B and the designed variants, the lanes show the insoluble fraction, soluble fraction, and purified protein, respectively. For commercial βLG, only the purified protein sample was analyzed.

**Figure S4.**
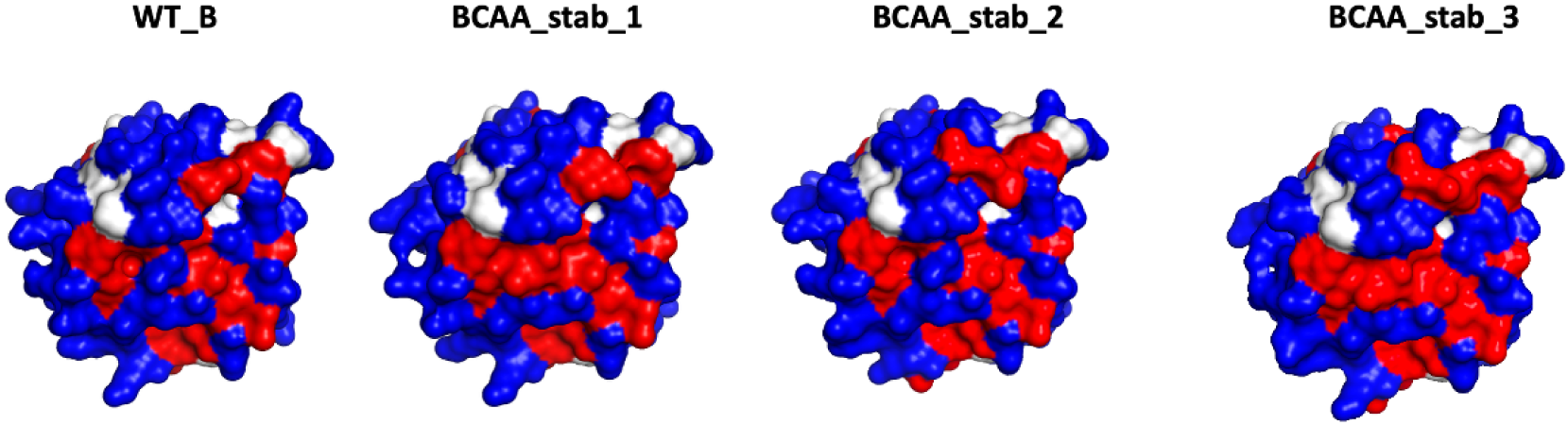
Predicted surface hydrophobicity of WT_B, BCAA_stab_1, BCAA_stab_2, BCAA_stab_3, BCAA_stab_4. Molecular surfaces were visualized in PyMOL using predicted protein structures. Hydrophobic residues (Ala, Val, Leu, Ile, Met, Phe, and Trp) are shown in red; polar and charged residues (Ser, Thr, Asn, Gln, Asp, Glu, Lys, Arg, and His) in blue; and intermediate residues (Gly, Pro, Tyr, and Cys) in white.

